# Anaerobic time-resolved serial crystallography captures CO dissociation and rebinding in oxygen-sensitive [FeFe]-hydrogenase

**DOI:** 10.64898/2026.07.22.740055

**Authors:** Ivan Voloshyn, Robin Stipp, Lukas Grunewald, Laura A. Kiesewetter, Samuel L. Rose, Shibom Basu, Cecilia Casadei, Monika Bjelčić, Daniele de Sanctis, Sebastian Westenhoff, Moritz Senger

## Abstract

Redox-active, oxygen-sensitive metalloenzymes catalyze key reactions in biological energy conversion and small-molecule activation. Understanding their mechanisms requires structural characterization of transient catalytic intermediates. Time-resolved serial crystallography (TR-SX) enables direct visualization of protein dynamics during catalysis under near-physiological conditions. Its application to oxygen-sensitive enzymes has remained challenging because strict anaerobic conditions must be maintained throughout sample preparation and data collection. Here, we establish an anaerobic room-temperature serial crystallography workflow for the oxygen-sensitive [FeFe]-hydrogenase *Cp*I and demonstrate its applicability by determining a room-temperature structure of the CO-inhibited H_ox_-CO state and following inhibitory CO dissociation and rebinding on the millisecond timescale via TR-SX. The presented room-temperature H_ox_-CO structure closely resembles previous cryogenic models and shows no detectable evidence of oxygen-induced degradation or significant radiation damage. Time-resolved measurements reveal no detectable structural rearrangements accompanying CO dissociation and rebinding beyond displacement and return of the inhibitory ligand, indicating a rigid catalytic architecture that may facilitate rapid catalysis. The presented workflow enables time-resolved structural studies of oxygen-sensitive metalloenzymes under physiologically relevant conditions and opens the way to direct visualization of catalytic intermediates in redox enzymes.

## Introduction

Redox-active metalloenzymes catalyze many of nature’s most fundamental energy-conversion and small-molecule activation reactions, including biological nitrogen (N_2_) fixation, carbon dioxide (CO_2_) fixation, and hydrogen (H_2_) production^1^. Despite their exceptional catalytic efficiencies and broad biotechnological potential, the molecular mechanisms underlying these transformations remain incompletely understood, as many catalytically relevant intermediates are transient, short-lived, and therefore remain inaccessible to conventional structural approaches.

Among these oxygen-sensitive metalloenzymes, [FeFe]-hydrogenases are of particular interest as they catalyze H_2_ production at turnover frequencies approaching 20,000 H₂ molecules per second^2,3^ at ambient temperatures and with small overpotentials^3–5^. These exceptional catalytic properties motivate their application in sustainable hydrogen production technologies^6,7^. Catalysis occurs at the H-cluster, a unique organometallic cofactor consisting of a [4Fe4S] cubane linked through a cysteinyl thiolate to a diiron subcluster coordinated by several CO and CN⁻ ligands as well as an azadithiolate (ADT) ligand (Fig. 1B). During catalysis, electrons are transferred to the H-cluster through a relay of iron-sulfur clusters while protons are delivered to the ADT ligand through a dedicated proton-transfer pathway (Fig. 1A). A terminally bound hydride ion (H^-^) is proposed to form at the open coordination site (OCS) of the distal iron (Fe_d_) of the H-cluster (Fig. 1B). Through protonation of this hydride, a H_2_ molecule is generated and released through hydrophobic gas channels^8^.

**Figure 1:**
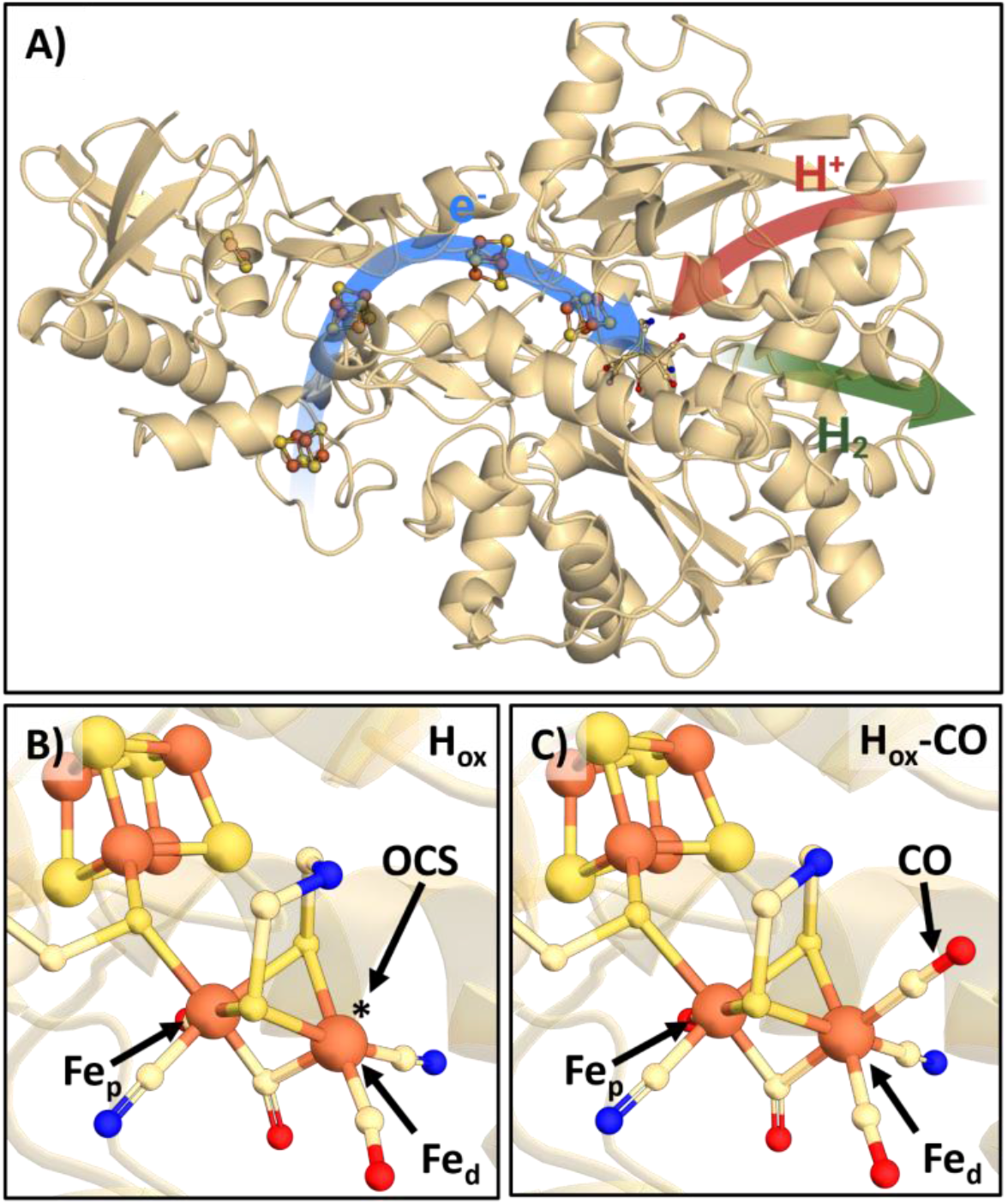
Structural representation of *Cp*I and the H-cluster; **A)** Overall structure of *Cp*I. Electrons are delivered to the H-cluster via the electron transfer chain (blue arrow), while protons are supplied by the proton transfer pathway (red arrow). After hydrogen formation, the molecular H_2_ leaves the enzyme through gas channels (green arrow). **B)-C)** Close up view of the H-cluster in the core of *Cp*I. In the H_ox_ state (**B)**, PDB code: 4XDC), the distal iron (Fe_d_) has an open coordination site (OCS). This site is occupied by an additional CO ligand in the H_ox_-CO state (**C)**, PDB code: 8ALN) of *Cp*I. Carbon atoms are shown in beige, sulfur atoms in yellow, oxygen atoms in red, nitrogen atoms in blue and iron atoms in orange.

The current understanding of this catalytic cycle was largely derived spectroscopically. Infrared (IR) spectroscopy exploits the CO and CN⁻ ligands of the H-cluster as intrinsic probes to monitor changes in the electronic structure during catalysis^9,10^. Using this methodology, several redox states of [FeFe]-hydrogenases were identified, including the oxidized ground state (H_ox_)^11^, the one-electron reduced intermediates (H_red_’, H_red_)^12–14^, two electron reduced intermediates (H_sred_^15^, H_hyd_^16–18)^, as well as the CO-inhibited state (H_ox_-CO, Fig. 1C)^19,20^. Time-resolved IR measurements have further resolved the kinetics of CO photodissociation and rebinding^21–23^. However, while spectroscopy has provided detailed information on the kinetics and electronic properties of catalytic states, it cannot directly reveal their structures.

X-ray crystallography has provided an important complementary view of the enzyme. Early crystal structures established the overall architecture of the H-cluster and proton-transfer pathway^24–27^, while more recent X-ray free-electron laser (XFEL) studies resolved the H_ox_ state free of radiation-induced artifacts at near-atomic resolution^28^. Despite these advances, all available [FeFe]-hydrogenase structures have been determined under non-physiological conditions. While cryocooling reduces radiation damage and enables complete data collection from single crystals, it may suppress conformational dynamics and introduce structural artifacts, especially after addition of cryo-protectants^29,30^. Consistent with this, spectroscopic studies have suggested temperature-dependent structural differences among catalytically relevant intermediates^24–26^. Whether existing structural models therefore accurately represent the conformational landscape of the active enzyme remains unresolved.

Time-resolved serial crystallography (TR-SX) provides a powerful approach for overcoming this limitation by enabling direct visualization of protein structures under near-physiological conditions^27^, while at the same time granting access to the thus far inaccessible structure of transient catalytic intermediates.

However, extending TR-SX to [FeFe]-hydrogenases has remained a major experimental challenge. [FeFe]-hydrogenases are irreversibly inactivated by molecular oxygen^31,32^, and thus require strict anaerobic conditions throughout crystal preparation, sample handling, reaction initiation, and X-ray data collection. In addition, the H-cluster and its associated electron-transferring relay of iron-sulfur clusters are susceptible to X-ray-induced reduction, making careful control of radiation damage essential for structural investigations of catalytically relevant redox states. Although serial crystallography minimizes cumulative radiation damage by exposing each microcrystal only once, the combined requirements of anaerobic sample handling, synchronized reaction initiation, and room-temperature (RT) serial data collection have thus far prevented TR-SX studies of [FeFe]-hydrogenases. Recent methodological developments have begun to overcome these limitations. TR-SX has provided unprecedented insight into the mechanism of cytochrome *c* oxidase^33–35^. Furthermore, RT serial crystallography studies of photosystem II^36,37^ demonstrated that oxygen-sensitive metalloenzymes can be investigated under strictly anaerobic conditions while preserving biologically relevant redox states. Furthermore, the development of different sample-delivery platforms^38^ has enabled the transfer of protein crystals from preparation to X-ray interrogation without atmospheric exposure. Together with advances in serial data collection at synchrotron and XFEL sources, these advances make time-resolved structural studies of [FeFe]-hydrogenase metalloenzymes feasible.

A further requirement for TR-SX is a reaction that is synchronized throughout the crystal ensemble. In [FeFe]-hydrogenases, the CO-inhibited H_ox_-CO state provides an experimentally accessible reaction trigger to enable this. Photodissociation of the exogenous CO ligand generates a well-defined starting point from which ligand rebinding can be followed over time. Previous time-resolved IR studies established the kinetics of this process by spectroscopic means^21,22^. Despite this spectroscopic characterization, the accompanying structural changes have remained unknown.

Here, we establish an anaerobic room-temperature serial crystallography workflow for oxygen-sensitive metalloenzymes using the [FeFe]-hydrogenase *Clostridium pasteurianum* hydrogenase I (*Cp*I) as a model system. We determine the first room-temperature structure of an [FeFe]-hydrogenase and combine synchrotron serial crystallography with *in crystallo* infrared spectroscopy to visualize inhibitory CO dissociation and rebinding on the millisecond timescale. Together, these results provide a framework for future studies of oxygen-sensitive redox metalloenzymes, opening a structural dynamics field for these enzymes.

## Results

### Establishing an anaerobic RT serial crystallography workflow for *Cp*I

While high-resolution crystal structures of the [FeFe]-hydrogenase *Cp*I have previously been reported for both the oxidized resting state, H_ox_ and the CO-inhibited state H_ox_-CO^39,40^, all available structures were determined at non-physiological temperatures. To enable structural studies at RT conditions, we established a serial crystallography workflow comprising optimized microcrystal production together with an anaerobic sample-delivery strategy that preserves the oxygen-sensitive H-cluster throughout sample handling and data collection.

The first requirement for serial crystallography was the generation of a reliable supply of well-diffracting microcrystals. By incorporating an additional seeding step into previously established crystallization protocols^23,41^, we reliably obtained microcrystals with typical dimensions of approximately 25 x 4 x 4 µm, well-suited for serial diffraction experiments (SI Fig. 2). *In crystallo* FTIR spectroscopy confirmed that CO treatment generated the expected H_ox_-CO state population prior to data collection (SI Fig. 4).

**Figure 2:**
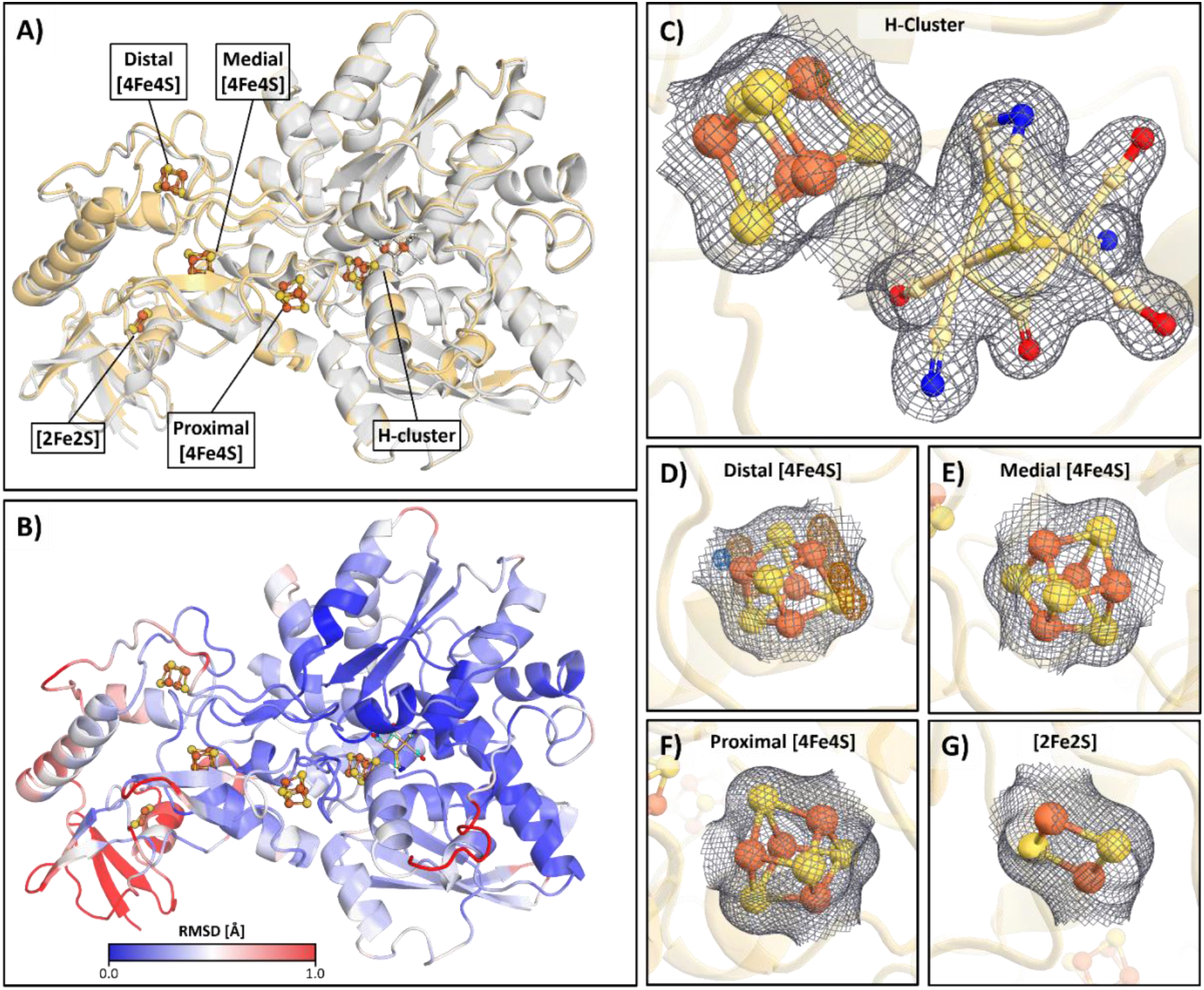
RT structure of *Cp*I shows strong overlap with cryogenic structure while displaying minimal radiation damage; **A)** RT H_ox_-CO structure in beige overlaid with a previously published cryogenic structure of *Cp*I (PDB code: 8ALN) in grey. **B)** RMSD deviation between RT and cryogenic H_ox_-CO structure shown above. RMSD is displayed via a blue to red gradient. **C)-G)** The RT dark structure is shown overlaid with the electron density map (grey mesh) at 1.5 σ and carved around the [4Fe4S], [2Fe2S] and H-clusters. In addition, the Fo-Fc difference electron density map is shown at 3.0 σ. Positive densities are shown in orange, negative densities in blue. Shown are the following FeS clusters: H-cluster **(C)**, distal [4Fe4S] cluster **(D)**, medial [4Fe4S] cluster **(E)**, proximal [4Fe4S] cluster **(F)** and [2Fe2S] cluster **(G)**.

Preserving the oxygen-sensitive H-cluster and accessory iron-sulfur cofactors during RT serial crystallography required an anaerobic sample-delivery strategy. Evaluation of several sheet-on-sheet (SOS) sample-delivery systems showed that SerialX fixed-target chips did not maintain anaerobic conditions, owing to sealing limitations and oxygen ingress during data collection. In contrast, both the foil fixed target setup at ID29 (ESRF)^42^ and the SOS-OS used at MicroMAX (MAX IV), each employing EVAL™ EVOH EF-F resin film, could be assembled reproducibly under anaerobic conditions and preserved the oxygen-sensitive H-cluster for more than one hour after transfer to ambient atmosphere, as verified by crystallography. These optimized sample-delivery strategies enabled reliable RT serial crystallography of the highly oxygen-sensitive [FeFe]-hydrogenase *Cp*I and provided the experimental foundation for the structural investigations described below.

### RT serial crystallography reveals a near-atomic resolution structure of *Cp*I

Using the anaerobic RT serial crystallography workflow established above, we determined the first RT (298 K) crystal structure of the [FeFe]-hydrogenase *Cp*I at the MicroMAX beamline at MAX IV. The resulting dataset yielded a 1.9 Å resolution structure of *Cp*I in the CO-inhibited H_ox_-CO state, providing the first near-atomic resolution view of this highly oxygen-sensitive enzyme under near-physiological temperatures.

The crystals displayed the P2₁ space group with two protein molecules in the asymmetric unit and four molecules per unit cell, consistent with previously reported cryogenic structures. This conserved crystal packing permits direct comparison with the cryogenic H_ox_-CO model (PDB: 8ALN)^43^ using least-squares superposition. The two structures exhibit a high degree of structural similarity, with an average root-mean-square deviation (RMSD) of only 0.446 Å (Fig. 2A), indicating that the overall protein architecture remains highly conserved at RT. Structural differences are largely confined to solvent-exposed surface regions, whereas the catalytic core, including the H-cluster, the surrounding second coordination sphere, and the proton transfer pathway, remain highly conserved (Fig. 2B). Within the active-site environment, only Met353 and Ser357 were modeled in alternate conformations. Interestingly, these are the same residues previously reported to exhibit conformational heterogeneity in synchrotron-derived cryogenic structures and proposed to be affected by X-ray-induced photoreduction^28^.

As expected, the RT structure exhibits higher overall atomic displacement than the cryogenic model, reflected by elevated B-factors (SI Fig. 3A-B). However, normalization of the B-factors reveals a highly similar pattern of relative flexibility between the two structures with only subtle differences (SI Fig. 3C-D), indicating that the increased mobility primarily reflects enhanced thermal motion rather than localized structural destabilization. Consistent with this interpretation, the RT structure displayed a modest expansion of the unit cell relative to the cryogenic structure (SI Table 1). Together, these observations demonstrate that the RT structure preserves the overall structure of *Cp*I displayed in previous cryogenic structures while allowing the increased thermal motion characteristic for physiological temperatures.

**Figure 3:**
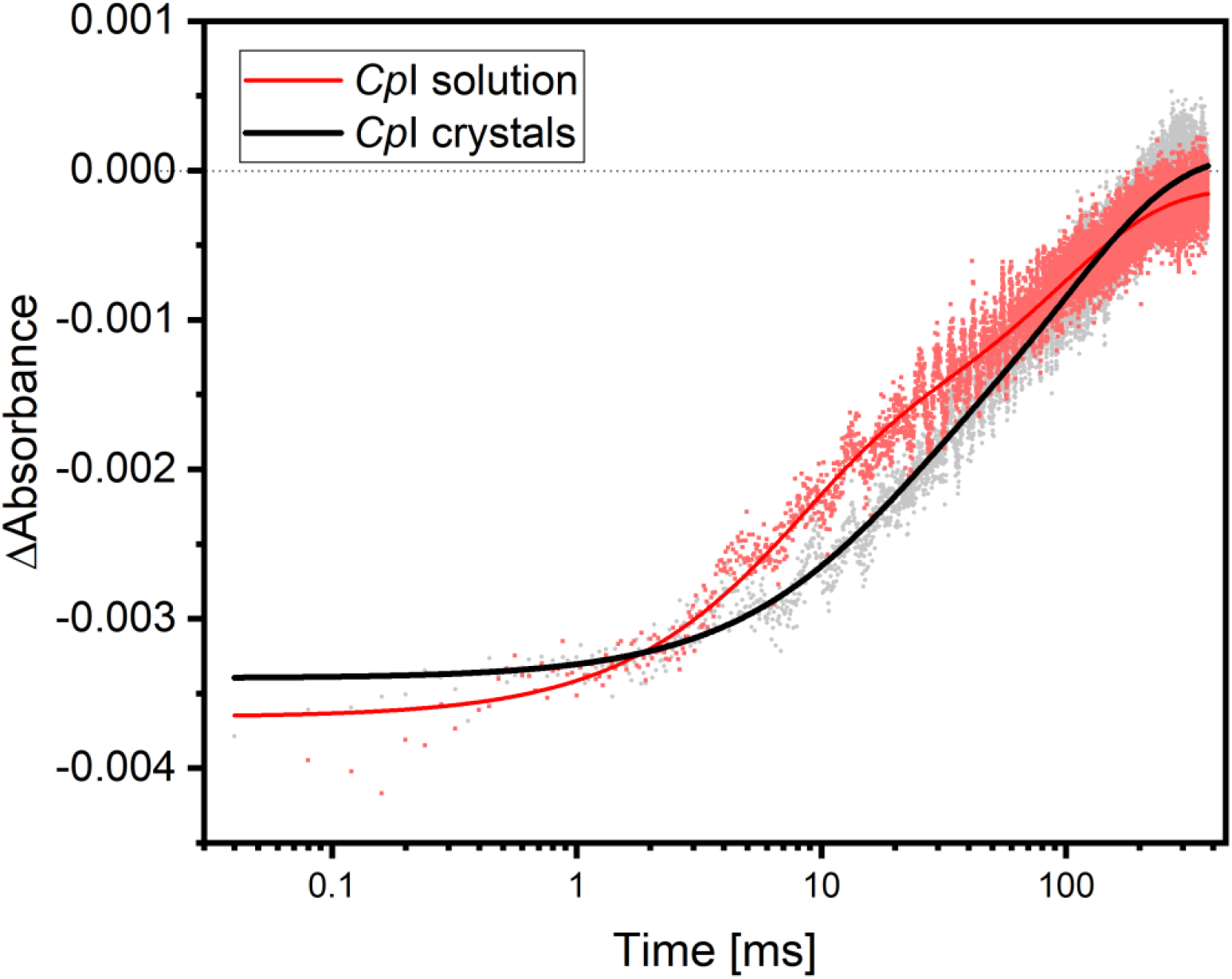
Infrared kinetics of CO rebinding in *Cp*I solution and crystals after photolysis; Difference absorbance changes at 2015 cm^-1^, corresponding to the coupled symmetric stretching vibration of the equatorial CO ligands of the H-cluster in the H_ox_-CO state. The changes were monitored from 50 ns to 300 ms after excitation with a 355 nm laser pulse. Measurements were performed for *Cp*I in solution (red) and in microcrystals (black). Solid lines represent bi-exponential fits to the experimental data. Kinetics were corrected for non-specific baseline changes by subtraction of the signal recorded at 2000 cm^-1^.

A key concern for RT serial crystallography at synchrotron sources is radiation damage, particularly for metalloproteins where metal centers and FeS clusters are among the most susceptible sites for X-ray-induced damage. Inspection of the five FeS clusters in *Cp*I structure reveals no significant difference density around the iron atoms at the proximal and medial [4Fe4S] cluster (Fig. 2E-F) or the [2Fe2S] cluster (Fig. 2G). Likewise, the geometry of the H-cluster closely matches previously reported cryogenic and XFEL structures of [FeFe]-hydrogenases (Fig. 2C), providing no evidence for either X-ray-induced structural perturbation or oxidative degradation. The only notable deviation is observed at the distal [4Fe4S] cluster (Fig. 2D). While the geometry of this metal cluster is consistent with previously reported models, several regions of positive difference density are present.

However, these features are spatially separated from both the metal ions and their coordinating residues, arguing against site-specific radiation damage. Instead, because these features are confined to a solvent-exposed surface region, that also exhibits elevated atomic mobility (SI Fig. 3), they most likely reflect increased conformational heterogeneity of the surrounding protein matrix. Metal-site occupancies close to unity confirmed complete retention of the iron-sulfur cofactors, indicating that neither significant radiation damage nor oxidative degradation occurred during data collection.

### Time-resolved IR spectroscopy establishes CO rebinding kinetics in *Cp*I microcrystals

Photodissociation of the inhibitory CO ligand from the H_ox_-CO state using 300-600 nm UV-Vis light provides a synchronized trigger for time-resolved experiments in [FeFe]-hydrogenases^21,22,44^. To define suitable time points for structural measurements, the kinetics of CO rebinding first needed to be characterized in *Cp*I microcrystals.

Time-resolved infrared (TR-IR) spectroscopy was used to follow dissociation and subsequent rebinding of the CO ligand over a time window spanning 50 ns to 300 ms following excitation with a 355 nm laser pulse. CO rebinding was monitored via the 2015 cm^-1^ absorption band, corresponding to the coupled symmetric stretching vibration of the equatorial CO ligands of the H-cluster^20^, which serves as a sensitive marker of the H_ox_-CO state^21,23^. To correct for non-specific baseline changes, including potential thermal effects, the kinetics were referenced to the signal at 2000 cm^-1^ (SI Fig. 5), where no cofactor-specific absorbance changes are expected within the investigated time window. Any potential vibrational excited states contributing to this region decay on the picosecond timescale ^22^, preceding the time resolution of our present measurement.

These baseline-corrected kinetics reveal CO rebinding with bi-exponential characteristics in both solution and crystalline samples (Fig. 3). In solution, the fast and slow phases exhibit time constants of 7.3 ± 0.2 ms and 93.6 ± 1.1 ms, respectively. In crystals, the corresponding time constants increase to 14.9 ± 0.7 ms and 107.4 ± 1.8 ms, indicating that the dominant fast rebinding process is approximately twofold slower in the crystalline state. The observed fast phase is consistent with previously reported CO rebinding kinetics for the [FeFe]-hydrogenase HydA1 from *Chlamydomonas reinhardtii* in solution with reported time constants ranging from 1 ms^23^, 3.9 ms^22^ to 13 ms^21^. The slow time constant contribution has been reported for HydA1^23^, although its contribution appears more pronounced in *Cp*I solution samples.

The slower rebinding observed in crystals indicates that the crystalline environment modestly alters ligand dynamics while preserving the overall photochemical reaction. Importantly, these measurements establish that photodissociation and subsequent CO rebinding can be reproducibly initiated and monitored in *Cp*I microcrystals under conditions compatible with time-resolved serial crystallography. The experimentally determined rebinding kinetics thereby define the time points for the structural investigation presented below.

### Time-resolved synchrotron serial crystallography captures CO rebinding in *Cp*I [FeFe]-hydrogenase

To directly visualize CO rebinding using TR-SX, we recorded diffraction data at the ID29 beamline (ESRF). *Cp*I microcrystals were embedded in a viscous hydroxy-ethyl-cellulose (HEC) matrix to inhibit migration of the crystals during measurement and applied to foil fixed-target support sealed with 12.5 µm EVAL™ EVOH EF-F film. Photolysis with a 375 nm diode triggered dissociation of the inhibitory CO ligand from the OCS at the distal iron of the H-cluster. Based on TR-IR data, four time points were selected to capture the early phase of CO rebinding: 3 ms, 7 ms, 11 ms and 15 ms (SI Fig. 1).

Difference electron density (DED) maps were calculated relative to the dark-state H_ox_-CO structure. The initial fluence of the pump diode at 1.8 mJ/mm^2^ was insufficient to yield any significant difference signal, leading us to increase the fluence to 5.4 mJ/mm^2^. Because the available excitation wavelength provided limited efficiency for CO photodissociation at the H-cluster and excitation of the iron-sulfur clusters contributed additional background signal, this relatively high pump fluence was required. This resulted in comparatively noisy DED maps, requiring careful assessment of the resulting difference densities. While the 3 ms and 7 ms datasets exhibited sufficient signal quality for direct interpretation, the signal in the 11 and 15 ms datasets was only slightly above the noise. As a result, these datasets were merged for subsequent analysis to increase signal-to-noise ratio.

The resulting DED maps revealed pronounced negative difference density at the position of the inhibitory CO ligand bound to the distal iron of the H-cluster in chain A at 3 ms after photoexcitation (Fig. 4A). This negative density decreases at 7 ms (Fig. 4B) and is weakest in the merged 11-15 ms dataset (Fig. 4C), consistent with progressive rebinding of the inhibitory CO ligand. Quantification of the integrated negative difference density within a 1 Å radius around the CO binding site confirms this gradual recovery of electron density over time (Fig. 4D). In an effort to decrease the influence of noise on data quality, METEOR denoising was explored and produced results consistent with raw data signals (SI Fig. 5), strengthening the abovementioned data.

**Figure 4:**
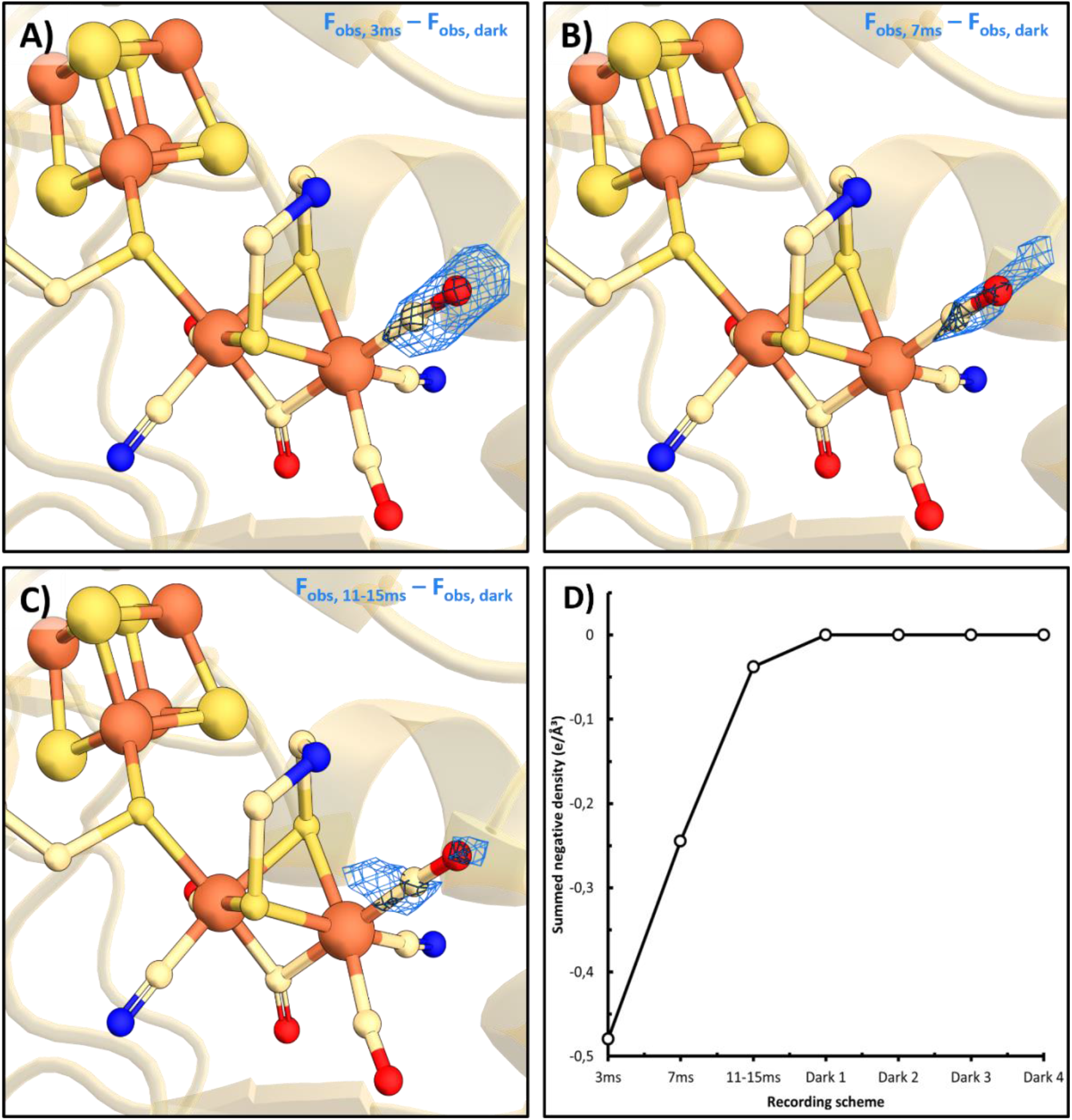
Rebinding of the inhibitory CO ligand after photolysis revealed by TR-SX. **A-C)** Fo−Fo DED map at 3, 7 and 11-15 ms showing negative density (blue mesh) around the inhibitory CO ligand. The DED is shown at 3 σ and carved around the ligand. **D)** Time-dependent decrease in summed negative density around the CO site in chain A (1 Å radius, −2 σ threshold), consistent with spectroscopic data (Fig. 3). All dark datasets lacked detectable negative DED features at the CO position.

Like chain A, negative density consistent with CO dissociation was also observed in chain B. However, no reproducible time-dependent trend could be established. This may reflect differences in excitation efficiency or local crystal environment between the two molecules in the asymmetric unit. Dark datasets collected between repeated illuminated measurements (SI Fig. 1) showed no negative difference density at the CO position, indicating that light contamination during data collection could be excluded (Fig. 4D).

Remarkably, no additional significant structural changes were detected beyond the inhibitory CO ligand itself. Neither the geometry of the H-cluster nor residues lining the gas channel or second coordination sphere exhibited significant conformational changes throughout the investigated time window. This observation indicates that CO rebinding occurs without detectable rearrangement of the surrounding catalytic environment on the millisecond timescale. Despite clear evidence for CO dissociation from the active site, we could not detect localized positive difference density corresponding to a transiently trapped CO molecule.

To further quantify the extent of ligand dissociation, the 3 ms, 7 ms and 11-15 ms structures were fitted using extrapolated structure factors (see Methods and SI Fig. 6), allowing estimation of the CO occupancy within the photoactivated crystal subpopulation. These refinements indicate near-complete photodissociation of the inhibitory CO ligand in this subset at 3 ms, with a refined occupancy of 3 %. Subsequently, the CO molecule gradually rebinds, reaching occupancies of 17 % at 7 ms and 29 % at 11-15 ms. Overlaying these estimated CO occupancies with our *in crystallo* TR-IR spectroscopy data (SI Fig. 8) provides additional evidence for successful crystallographic capture of CO photodissociation and rebinding in *Cp*I microcrystals by TR serial crystallography.

## Discussion

This work establishes RT serial crystallography as a viable approach for structural studies of the oxygen-sensitive [FeFe]-hydrogenase *Cp*I and provides the first near-atomic resolution view of the enzyme under physiologically relevant conditions.

Compared to the previously published cryogenic structures, the catalytic core surrounding the H-cluster remains remarkably well conserved at RT. Subtle temperature-dependent differences are observed in solvent-exposed region, particularly near the auxiliary [2Fe-2S] cluster, highlighting increased flexibility in peripheral regions of the enzyme. The close agreement between our RT structure and previous cryogenic models is particularly noteworthy in light of spectroscopic studies that suggested a rotated active-site geometry with a blocked open coordination site in the reduced diiron site intermediate H_red_ of [FeFe]-hydrogenases at RT^26,45,46^. Instead, our findings are consistent with growing evidence that a conserved active-site geometry for the reduced diiron site intermediate can be observed at RT^47^.

Using inhibitory CO photodissociation as an optical trigger, our time-resolved serial crystallography study of *Cp*I shows that dynamics can be monitored structurally in [FeFe]-hydrogenase crystals. TR-IR spectroscopy revealed modestly slower CO rebinding in crystals compared to solution, consistent with restricted lattice dynamics. TR-SX measurements proved highly compatible with this data in the millisecond range. A striking finding of our study is that CO dissociation and rebinding occur without detectable structural rearrangements beyond loss and rebinding of the inhibitory ligand. The H-cluster and surrounding gas-channel residues remain conformationally unchanged within the observed time window. These results suggest that the *Cp*I catalytic core remains structurally rigid during ligand exchange, allowing CO to diffuse through the gas channel without the need to form structurally altered intermediate states. The apparent structural preorganization and high rigidity of the active site may contribute to the exceptionally high turnover rates of [FeFe]-hydrogenases, as extensive structural rearrangements during catalysis would be incompatible with fast H_2_ turnover. We previously showed spectroscopically that only the hydrogen bonding network of the proton transfer pathway rearranges and excluded structural changes when transferring from the oxidized H_ox_ state to the reduced diiron site state H_red_ in fast [FeFe]-hydrogenase *Cr*HydA1 from *Chlamydomonas reinhardtii*^48^. In contrast in the slow catalysis, potentially sensing [FeFe]-hydrogenase *Tam*HydS from *Thermoanaerobacter mathranii* we could detect secondary structure changes which accompany the H_ox_ to H_red_ transition^49^. This is consistent with the structural evidence derived from our structures and paints a picture of finely modulated rigidity as one measure of tuning the respective [FeFe]-hydrogenases characteristics dependent on their role.

Our TR-IR spectroscopy revealed baseline shifts consistent with transient heating of the microcrystals upon illumination, an effect that was minimal in solution and likely enhanced by the high protein concentration and PEG/glycerol-rich crystallization medium. High light fluences were required for efficient photolysis, resulting in increased thermal motion and elevated noise in the difference electron density maps. Nevertheless, clear CO dissociation and rebinding signals were observed. However, CO photodissociation is initiated on the picosecond timescale, highlighting the need for time-resolved XFEL experiments to capture the earliest structural events following photoexcitation. Serial XFEL-based measurements would also avoid synchrotron-induced photoreduction effects affecting residues such as Met353 and Ser357, providing a more faithful view of the RT enzyme. Beyond inhibitory CO photolysis, the anaerobic RT serial crystallography framework established here opens the possibility of visualizing the catalytic cycle of [FeFe]-hydrogenases under near-physiological conditions. Photo excitable electron donors can trigger electron transfer to the H-cluster, enabling light-driven catalysis in this otherwise non-photoactive enzyme^48–55^. Coupling these artificial photo activation strategies with TR-SX provides a novel route to directly capture the structure of catalytic intermediates and extends the applicability TR-SX to solve the catalytic mechanism of oxygen-sensitive redox-active metalloenzymes with applications in biotechnology and beyond.

## Author contributions

Conceptualization: MS Methodology: IV, RS, MS

Investigation: IV, RS, LG, LAK, SLR, SB, MB, CC, DdS, MS

Visualization: RS, LG

Supervision: MS, SW

Writing - original draft: RS, IV, MS Writing - review and editing: all authors

## Acknowledgements

The Novo Nordisk Foundation (Grant ref. NNF23OC0085682, M.S), the Swedish Research Council (VR, grant no 2023−04593, I.V. and M.S.), FORMAS (2024-00586_3, M.S.) and the Carl Tryggers Stiftelse (CTS 24:3568, R.S and M.S.) are gratefully acknowledged for funding.

We acknowledge the MAX IV Laboratory for beamtime on the MicroMAX beamline under proposal 20250720. Research conducted at MAX IV, a Swedish national user facility, is supported by Vetenskapsrådet (Swedish Research Council, VR) under contract 2018-07152, Vinnova (Swedish Governmental Agency for Innovation Systems) under contract 2018-04969 and Formas under contract 2019-02496. MicroMAX is funded by the Novo Nordisk Foundation under the grant number NNF17CC0030666

We acknowledge the European Synchrotron Radiation Facility (ESRF) for provision of synchrotron radiation facilities under proposal MX-2778 and on beamline ID29.

## Methods

### Protein expression and purification

[FeFe]-hydrogenase from *C. pasteurianum* containing a C-terminal *Strep*-II tag was recombinantly overexpressed in *E. coli* BL21 (DE3) D*iscR* as previously reported^56^ with minor adjustments. The cultures were incubated at 20 °C for 20 hours after induction. Cells were aerobically harvested and transferred to a Vigor anaerobic glovebox filled with Argon and resuspended in lysis buffer (100 mM Tris-HCl (pH 8.0), 150 mM NaCl, 1.2 mg/mL lysozyme, 0.06 mg/mL DNase I, 0.06 mg/mL RNase A, 2.4 mg/mL MgCl_2_ * 6H_2_O, one cOmplete^TM^ EDTA-free protease inhibitor cocktail per 50 mL of buffer, 5 mg/mL sodium deoxycholate and 5 mg/mL sucrose). Cells were pre-treated for 1 hour in lysis buffer before being sonicated at 4 °C for 5 cycles (15 seconds on, 30 seconds off). The sonicated lysate was ultracentrifuged, filtered and purified using ÄKTA^TM^ Go system with a StrepTrap^TM^ XT column (Cytiva). The column was washed with wash buffer (100 mM Tris-HCl (pH 8.0), 150 mM NaCl) and the protein was eluted with wash buffer supplemented with 50 mM biotin. The eluted *Cp*I apo-protein was concentrated to 100 mg/ml, flash-frozen and stored at -80 °C. Protein purity was analyzed with SDS-PAGE and concentration was measured using Pierce^TM^ Bradford Assay Kit (Thermo Scientific). [Fe] content was quantified using an Iron Assay Kit (MAK472, Sigma-Aldrich).

*Cp*I holo-enzyme was obtained as described before^57^ in the absence of sodium dithionite, followed by desalting with a HiPrep^TM^ 26/10 Desalting column equilibrated in 10 mM Tris-HCl (pH 8.0). The holo-enzyme was concentrated, aliquoted, flash-frozen and stored at -80 °C. Concentration and [Fe] content were measured for the purified protein using the previously mentioned kits. Cofactor integration and *Cp*I activity were verified using FTIR spectroscopy.

### Transmission FTIR spectroscopy

A solution of 4 µL of *Cp*I protein (73 mg/mL) in 10 mM Tris-HCl buffer pH 8 was deposited on the CaF_2_ window in the anaerobic Ar atmosphere of a glove box (Vigor). The sample was sealed with a second CaF_2_ window and mounted on a custom build transmission cell which allowed for gas exchange and illumination. The sealed cell was then purged with CO gas for at least 5 minutes. The cell was installed in a FTIR spectrometer (Vertex V70v, Bruker) and complete CO inhibition was confirmed recording spectra with 1 cm^-1^ resolution, a scanner velocity of 80 Hz and varying number of scans (typically 100 scans). All measurements were performed at ambient conditions (RT and atmospheric pressure). In case of crystals, 4 µl of high-concentration microcrystal slurry (crystallization conditions: 0.1 M MES pH 6.0, 0.4 M MgCl_2_, 20 % glycerol, 20 % PEG 4000) was applied and the same procedure was followed.

### Time resolved quantum Cascade Laser infrared spectroscopy

For RT (∼22 °C) transient absorption measurements, a Q-switched Nd:YAG laser (Ekspla) was employed to obtain 355 nm pump light with a ∼50 mJ per pulse and a FWHM of 10 ns. Probe light was provided by a continuous-wave IR quantum cascade laser (QCL) with a tuning capability between 1960 and 2150 cm^−1^ (Daylight Solutions). The IR probe light was superimposed on the laser beam in a quasi-collinear arrangement at an angle of ca. 25 °. For IR detection, a liquid-nitrogen-cooled mercury−cadmium−telluride (MCT) detector (KMPV10-1-J2, Kolmar Technologies, Inc.) was used. Transient absorption traces were acquired with a Tektronix TDS 3052B 500 MHz (5GS/s) oscilloscope in connection with the L900 software (Edinburgh Instruments) and processed using Origin 2019 software.

### Protein crystallization

*Cp*I crystals were produced using microbatch crystallization. For this, purified protein solution with a concentration of 20 mg/ml was mixed with reservoir buffer (0.1 M MES pH 6.0, 0.4 M MgCl_2_, 20 % glycerol, 20 % PEG 4000) in 1:1 (v:v) ratio and incubated on a tube rotator at 277 K. After 16 hours, rod-like brown macrocrystals (90 x 15 x 15 µm) formed. Crystal size was determined by applying the crystal slurry on a Burker-Turk cell counter followed by visual assessment under a light microscope.

To produce microcrystals suitable for serial crystallography, seeding was required. The microcrystal precipitation solution was prepared by crushing macrocrystals resuspended in their reservoir solution using a Seed Bead Steel^TM^ kit (Hampton Research). The suspension was diluted with reservoir buffer in 1:3 (*v*:*v*) ratio (crushed crystals: reservoir buffer). Subsequently, the resulting mixture was centrifugated at 1500 x g for 45 seconds to sediment large crystal shards and supernatant containing crystal seeds was taken.

*Cp*I microcrystals were prepared by mixing protein solution at 20 mg/mL with the microcrystal precipitation solution in 1:1 (*v*:*v*) ratio and incubating as described above. After 12 hours, rod-like microcrystals were formed (25 x 5 x 5 µm).

### CO inhibition of *Cp*I microcrystal for SSX

*Cp*I microcrystals were anaerobically harvested into PCR microtubes and sealed inside glass crimp-top headspace vials with the tube cap open. The sealed vials were injected with CO gas for 5 minutes using a needle at slight overpressure to prevent atmospheric oxygen from damaging the protein, followed by purging the vial with N_2_ gas. The purged vials were brought back inside anaerobic chamber, and the PCR microtubes were closed to prevent excessive evaporation of reservoir solution. The CO inhibition of *Cp*I microcrystals was confirmed by FTIR spectroscopy (SI Fig. 3).

### Data collection and analysis

In preparation of data collection, *Cp*I-H_ox_-CO microcrystals were mixed with 13 % hydroxy-ethyl-cellulose prepared with reservoir buffer in 30:70 (v:v) ratio inside a Coy anaerobic glovebox filled with 97.5 % N_2_ and 2.5 % H_2_ atmosphere. The microcrystals, now immobilized in viscous matrix, were applied onto a sheet-on-sheet (SOS) fixed-target chip^58^ and sealed anaerobically with a EVAL™ EVOH EF-F film. The sample application was performed under red light to minimize potential light-induced CO dissociation.

High-resolution diffraction data of the dark state *Cp*I-H_ox_-CO structure was collected at RT at the MicroMAX beamline at MAX IV with an X-ray beam size of 14 x 5 µm^2^ (FWHM), through a 15 µm diameter aperture, at a photon energy of 12.6 keV and flux of 2.78 x 10^12^ photons/s^59^, using SOS-OS chips^58^. The chip was raster scanned through the X-ray beam in a regular serpentine pattern defined by a mesh grid drawn defined in MXCuBE3^60^. The fast shutter was open during each row of the mesh. Rotation of the goniometer was blocked during data collection. Diffraction images were recorded by an EIGER2 9M CdTe (DECTRIS) detector with an exposure time of 10 ms. Sample viewing, alignment, and measurement were carried out using the beamline control software MXCuBE3^60^.

Time-resolved data collection of *Cp*I-H_ox_-CO microcrystals (dark and illuminated states) was performed at RT at ID29 beamline (European Synchrotron Radiation Facility). The dataset of CO-inhibited microcrystals was collected by performing a grid scan of SOS-OS^58^ chips using a 2 x 8 µm, 70 µs X-ray probe beam with vertical and horizontal spacing of 20 µm, at a photon energy of 11.56 keV and fluence of 1.2 x 10^15^ photons/s. CO dissociation from the H-cluster was triggered using a FlexxRay 375 nm pump diode light pulse with a 1 ms pulse duration, 80 µm (1/e^2^) with a fluence of 5.4 mJ/mm^2^, followed by four consecutive X-ray probes with a 4.3 ms delay between each X-ray pulse (SI Fig. 1). This resulted in four consecutive timepoints with a time delay of 3, 7.3, 11.6 and 15.9 ms respectively. Diffraction patterns were collected on a Jungfrau 4M detector. Hit finding was performed with Lima2 software.

### Processing of diffraction images

Indexing and integration of serial data was done with CrystFEL package^61^. Images were indexed using the xgandalf algorithm. Indexed patterns were then scaled and merged using partialator with the unity model and Friedel pair merging (point group = 2/m) to yield observed intensities |Iₒᵦₛ|. Comparehkl and checkhkl was used to compute the data quality statistics (See SI Table 2-3).

### Dark structure refinement

Molecular replacement with Phaser-MR was used to solve the initial phases using the existing cryo structure of *Cp*I (PDB ID: 8ALN) as a search model. Structure refinement was performed using PHENIX^62^ accompanied by manual model building steps with Coot^63^. The final structure had Rwork/Rfree of 0.15/0.19.

### Calculation of Difference Electron Density Maps

Difference electron density (DED) maps were calculated by Fourier transformation of the weighted difference of the observed structure factor amplitudes:

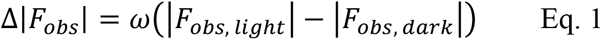

where phases were taken from the refined dark-state model. The weighting factor *ω* was applied individually to each reflection (hkl) to reduce the influence of outliers and was calculated according to the procedure described by Ursby and Bourgeois^64^

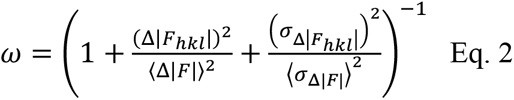

The weighting scheme depends on the magnitude of the individual difference structure factor Δ|*F*_ℎ*kl*_| and its associated uncertainty 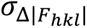 relative to the average values (Δ|*F*|, *σ*_Δ|*F*|_) across the dataset.

The Fourier transformation was performed in CCP4 using the FFT program. Prior to subtraction the observed structure factor amplitudes were scaled using SCALEIT in CCP4. First, |*F_obs_*_,*dark*_ | was scaled against the refined dark-state model|*F_calc_*_,*dark*_ |. Afterwards, |*F_obs_*_,*lig*ℎ*t*_| was scaled to the adjusted |*F_obs_*_,*dark*_ | dataset. This sequential scaling ensures that both datasets are on a common absolute scale before calculation of the DED maps. A detailed description of the calculation of DED maps is provided by Schmidt^65^.

### Light structure refinement

The light structures for 3, 7 and 11-15 ms were refined using extrapolated structure factors:

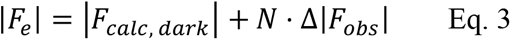

where |*F_calc_*_, *dark*_| are the calculated structure factors of the refined dark structure, Δ*F_obs_* are described above in Eq.1 and N is the extrapolation factor. N is linked to the photoexcitation yield through 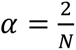, where the factor of 2 arises from the difference Fourier approximation described by Henderson and Moffat^66^. N was estimated by analyzing the negative intensity in the extrapolated maps as a function of N (SI Fig. 7). We estimated N to be approximately 7.14 %. In the first refinement step, the phases of the dark structure were used and resulted in very high Rwork/Rfree.

To reduce the Rwork/Rfree values to acceptable ranges, the structures were refined against phased corrected extrapolated maps. This procedure is described in detail by Schmidt as well (see Eq. 26)^65^.

