## Supplementary Information for "Anaerobic time-resolved serial crystallography captures CO dissociation and rebinding in oxygen-sensitive [FeFe]-hydrogenase"

| Cell Parameters | $H_{ox}$-CO (RT) | $H_{ox}$-CO (cryo) |
|---|---|---|
| a [Å] | 92.36 | 89.86 |
| b [Å] | 74.47 | 71.85 |
| c [Å] | 104.36 | 103.23 |
| a [°] | 90 | 90 |
| b [°] | 98.08 | 97.47 |
| g [°] | 90 | 90 |

**Supplementary Table 1: Comparison of cell parameters between RT and cryo structures of *Cp*I;** shown are the cell parameters of the structures of the $H_{ox}$-CO states of *Cp*I at room temperature (RT) or cryogenic temperatures. The PDB condes for the structure of the $H_{ox}$-CO (cryo) is 8ALN.

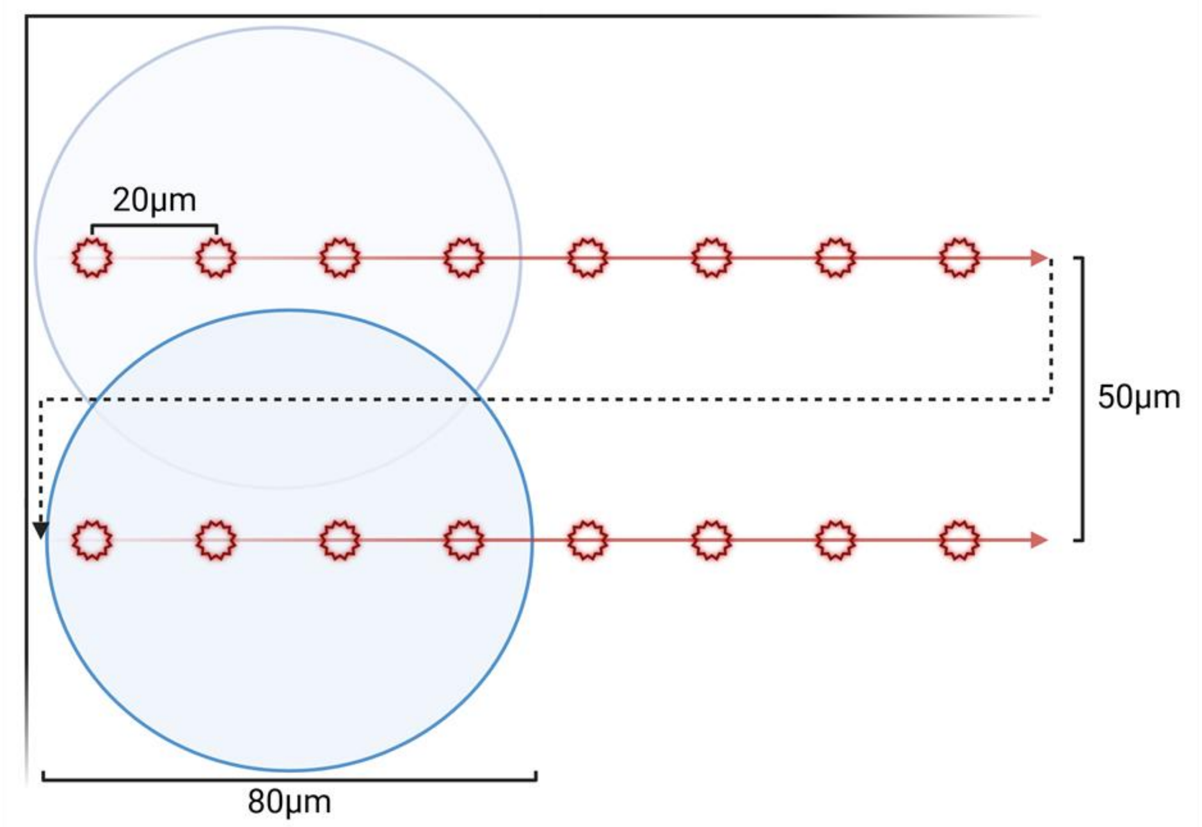


**Supplementary Figure 1: Representation of recording scheme for time-resolved data collection at ESRF.** A pump laser with a wavelength of 375 nm illuminates a circular area with 80 µm diameter (blue circle). 3 ms after the pump laser illumination, four consecutive x-ray pulses probed four spots (red stars) in the illuminated area along a linear path. Continuing the pattern, four dark data points were sampled. This pattern of eight probing cycles was repeated until a row of the chip was completely scanned. Afterwards, the sampling position returned to the left of the chip but shifted down by 50 µm to avoid sampling previously irradiated areas and exclude light contamination. The pattern was repeated until the chip was completely scanned.

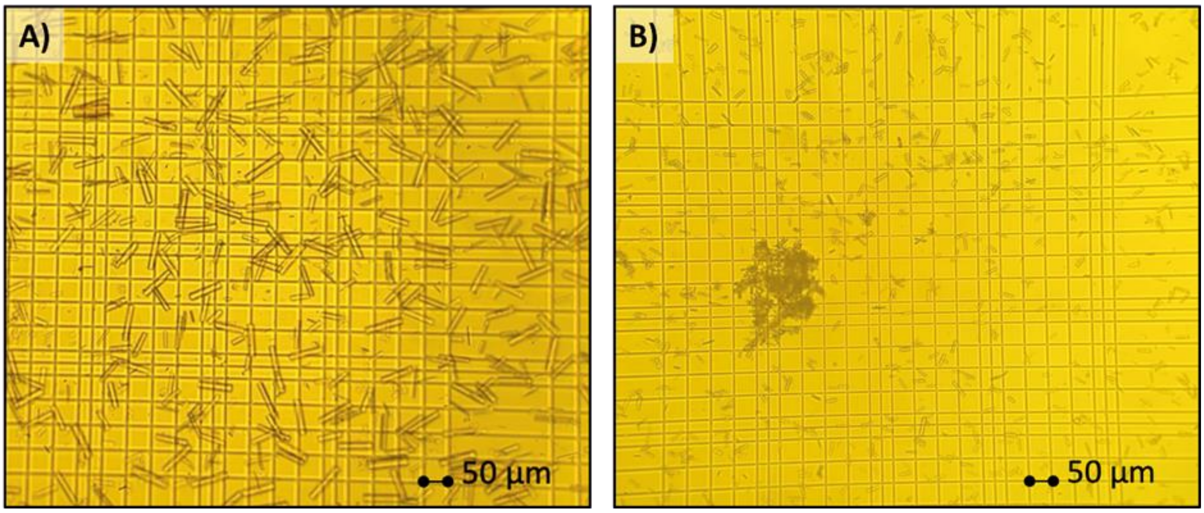


**Supplementary Figure 2: CpI crystals; A)** *Cp*I crystals generated following seed-less protocol. The crystals were subsequently crushed to generate seeds for subsequent crystallization batches. **B)** *Cp*I microcrystals grown with the addition of seeds during crystallization.

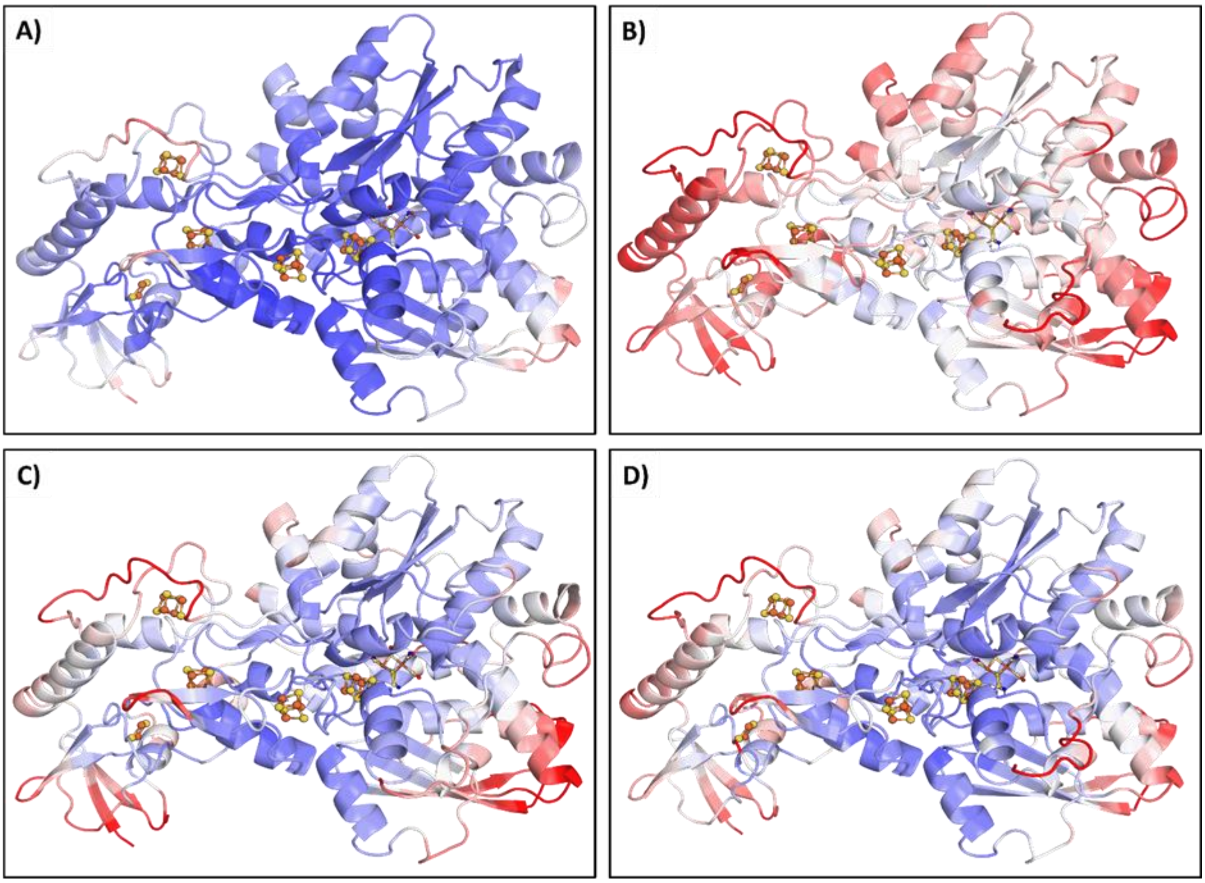


**Supplementary Figure 3: Comparison of B-factor distributions in cryogenic and RT structures of *Cp*I;** Cartoon representations are colored according to residue B-factors using a blue-white-red gradient, where blue indicates low B-factors and red indicates high B-factors. Shown are the cryogenic (**A**) and RT (**B**) structures colored by their refined B-factors. Panels (**C**) and (**D**) show the same structures colored by normalized B-factors, highlighting regions of relatively lower and higher flexibility within each structure independently.

**Supplementary Table 2: Statistics of dark datasets collected at MicroMAX and ESRF.**

| | **MicroMAX** | **ESRF** |
|---|---|---|
| number of indexed images | 212567 | 665041 |
| number of total reflections | 114284 | 346051 |
| possible reflections | 110075 | 62606 |
| Data completeness (%) | 100 (100) | 100 (100) |
| Overall CC1/2 | 0.99(0.68) | 0.99 (0.38) |
| Overall CC* | 0.99 (0.9) | 0.99 (0.74) |
| SNR | 9.19 (1.10) | 11.59 (1.54) |
| Redundancy | 1469.22 (1045.4) | 12942 (14801) |
| R-split (%) | 6.63 (101.3) | 5.99 (89.68) |
| Rwork/Rfree | 0.15/0.19 | - |
| Resolution | 39.84 - 1.90 | 74.47-2.30 |
| Refinement strategy | Classical | |
| Point group | 2 / m_uab | |
| Space group | P 1 21 1 | |
| Unit cell lengths and angles | 92.475 74.483 104.338 90 98.06 90 | 92.36 74.47 104.36 90 98.08 90 |

**Supplementary Table 3: Statistics of light datasets collected at ESRF.**

| | **3 ms** | **7 ms** | **11-15 ms** |
|---|---|---|---|
| number of indexed images | 175859 | 170577 | 351632 |
| number of total reflections | 340843 | 340738 | 343628 |
| possible reflections | 48827 | 48827 | 48827 |
| Data completeness (%) | 100 (100) | 100 (100) | 100 (100) |
| Overall CC1/2 | 0.99 (0.34) | 0.99 (0.29) | 0.99 (0.52) |
| Overall CC* | 0.99 (0.72) | 0.99 (0.67) | 0.99 (0.83) |
| SNR | 7.50 (1.36) | 7.37 (1.32) | 10.60 (1.88) |
| Redundancy | 3316 (3572) | 3219 (3467) | 6673 (7195) |
| R-split (%) | 10.07 (97.57) | 10.23 (103.55) | 7.20 (71.43) |
| Rwork/Rfree | 0.18/0.25 | 0.19/0.24 | 0.16/0.22 |
| Riso (to dark) | 0.101 | 0.102 | 0.089 |
| Resolution | 74.47-2.50 | | |
| Refinement strategy | Extrapolation | | |
| Photoactivation | ~7.14% | | |
| Point group | 2 / m_uab | | |
| Space group | P 1 21 1 | | |
| Unit cell lengths and angles | 92.475 74.483 104.338 90 98.06 90 | | |

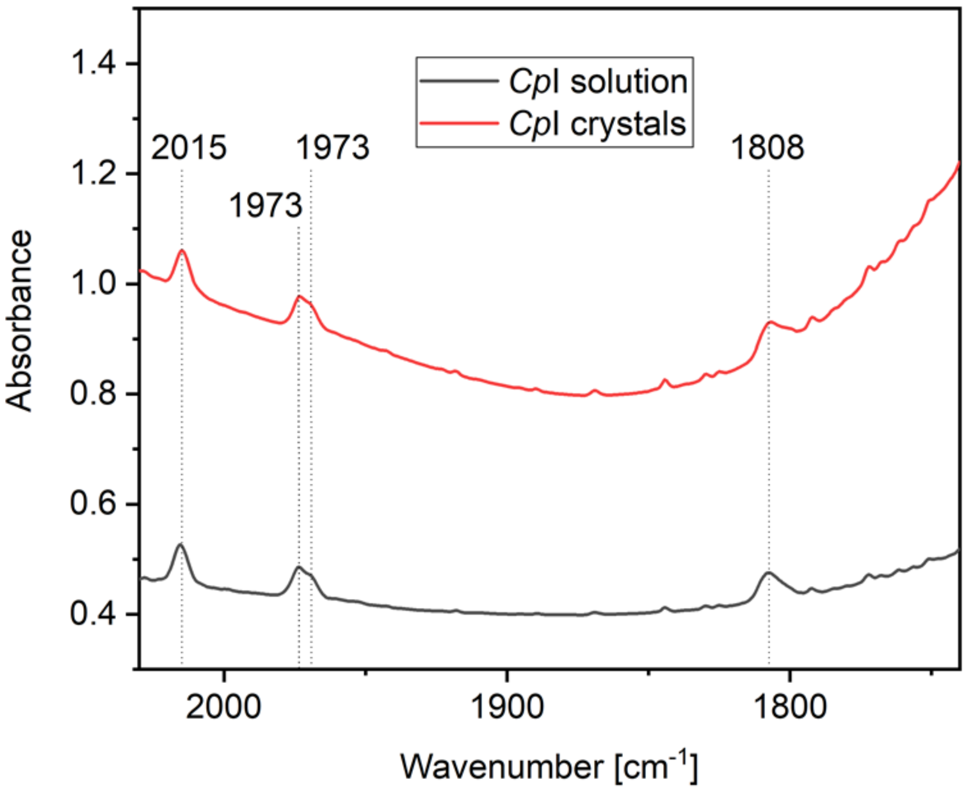


**Supplementary Figure 4: Fourier transform infrared transmission absorbance spectra of *Cp*I protein solution and crystal samples inhibited with CO.** Both *Cp*I samples (solution in black and crystals in red) show exclusively the characteristic CO ligand bands for $H_{ox}$-CO at 2015, 1973, 1969 and 1808 $cm^{-1}$, reporting on complete CO inhibition. Spectra are offset for clarity.

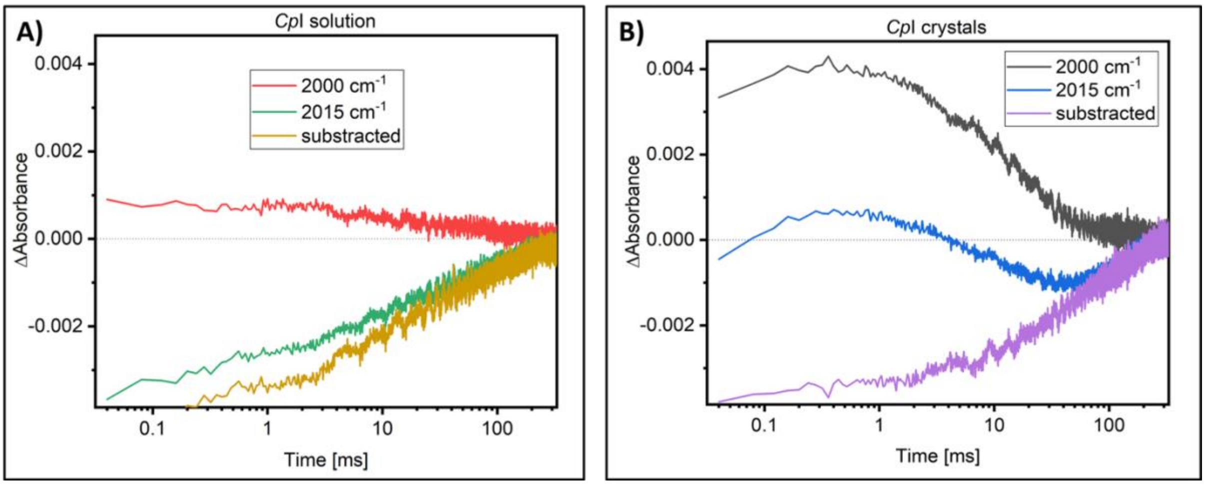


**Supplementary Figure 5: Transient Quantum Cascade Laser based infrared spectroscopy kinetics.** Raw kinetic traces recorded at 2000 $cm^{-1}$ (baseline drift) and 2015 $cm^{-1}$ (CO ligand band characteristic for $H_{ox}$-CO). Also show is 2015 $cm^{-1}$ kinetics with subtraction of the 2000 $cm^{-1}$ baseline signal reporting on the kinetics of CO rebinding only (as shown in Fig.3 main text). A: *Cp*I solution, B: *Cp*I crystals.

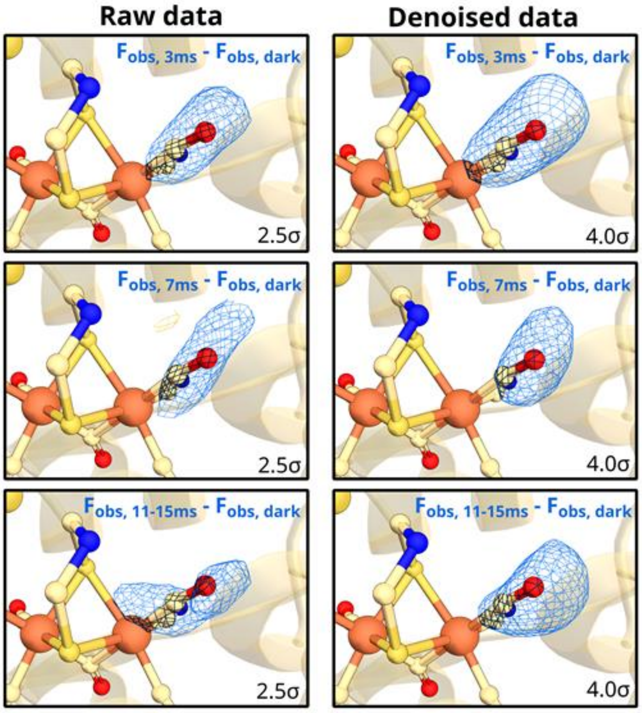


**Supplementary Figure 6: Comparison of raw and meteor-denoised DED maps for all recorded time points.** Meteor[67] reduces high-frequency noise and improves the clarity of the negative density surrounding the CO ligand. This also shows that the negative signal around the CO molecule is not noise.

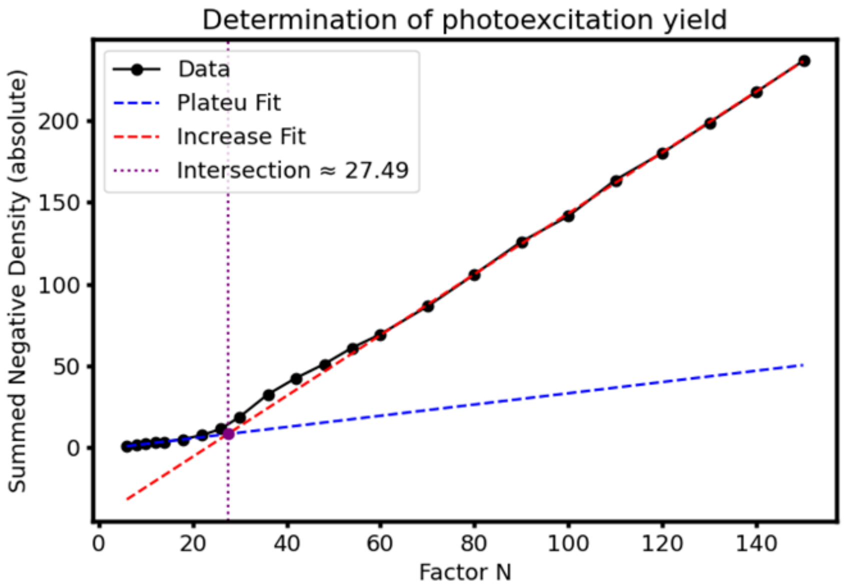


**Supplementary Figure 7**: **Determination of the photoexcitation yield of the 3 ms timepoint** using the sum of negative electron densities around the inhibitory CO molecule as a function of extrapolation factor N (see Methods). The extrapolation factor was determined to be ~28, which is equivalent to 7.14 % photoactivation yield. The extrapolation factor was assumed to be the same for the later timepoints.

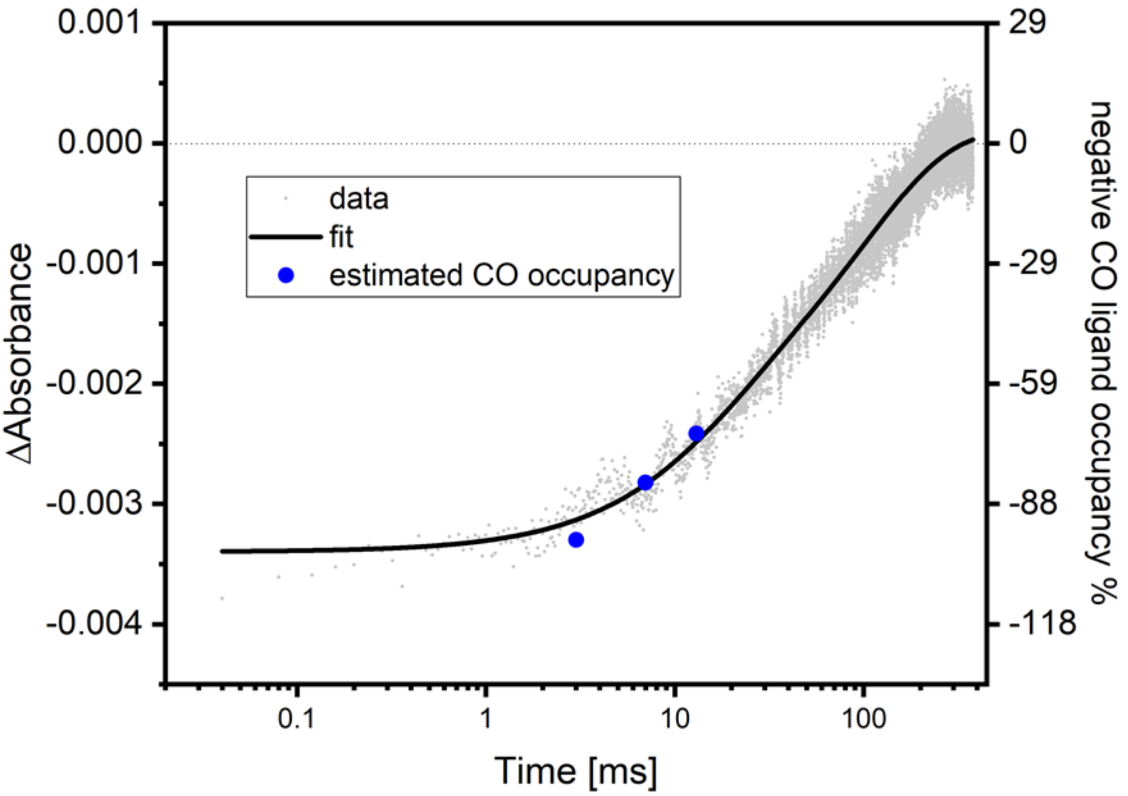


**Supplementary Figure 8**: **Overlay of TR-IR data and estimated CO ligand occupancies.** Difference absorbance changes at 2015 $cm^{-1}$, corresponding to the coupled symmetric stretching vibration of the equatorial CO ligands of the H-cluster in the $H_{ox}$-CO state. The changes were monitored from 50 ns to 300 ms after excitation with a 355 nm laser pulse. Data shown for *Cp*I in microcrystals (gray). Solid black line represents the biexponential fit to the experimental data. Kinetics were corrected for non-specific baseline changes by subtraction of the signal recorded at 2000 $cm^{-1}$. In blue dots show the estimated CO ligand occupancy in the photoactivated species of *Cp*I in our crystals.
